# Impact of gestational antibiotics on maternal and offspring gut microbiota and growth in pigs

**DOI:** 10.1101/2025.11.08.687112

**Authors:** Md. Rayhan Mahmud, Toomas Orro, Camilla Munsterhjelm, Tiina Pessa-Morikawa, Kristina Ahlqvist, Sami Junnikkala, Emilia König, Meritxell Pujolassos, M. Luz Calle, Huyk Nam Kwon, Claudio Oliviero, Mari Heinonen, Mikael Niku

## Abstract

Maternal microbiota modulates the development of the microbiota in the offspring. Effects of gestational antibiotics are not well understood, as most studies have focused on the perinatal period. We treated sows with penicillin, tetracycline or saline on days 78-80 of the 114-119-day gestation, a critical period in the fetal immune system development. Microbiotas were analyzed by 16S rRNA gene amplicon sequencing in sow feces and vagina at days 77 and 113, in colostrum, and in piglet feces at three days, three weeks and ten weeks of age. Sow fecal microbiota changed during pregnancy, but less in antibiotic groups. No significant effects on sow microbiota remained on day 113. The piglets of the antibiotic-treated sows had lower Firmicutes+Actinobacteriota to Bacteroidota+Proteobacteria ratio, lower alpha diversity and higher relative abundance of *Escherichia* at three days. At ten weeks, they exhibited higher alpha diversity, had higher of *Prevotella* and *Clostridium* sp*. CAG-127,* and smaller increase of *Limosilactobacillus* than the control. Increased alpha diversity at 10 weeks was associated with lower weight gain during nursing. *Oliverpabstia* and *Mitsuokella* were positively associated with growth, while *CAG-127* and *Campylobacter B* were negatively associated. Sow antibiotic treatment decreased the positive effect of *Oliverpabstia* and *Mitsuokella* and increased the negative effects of *CAG-127* and *Campylobacter B*. Gestational antibiotics may have had adverse effects on microbiota and growth of the offspring, even if their effects on maternal microbiota were undetectable by parturition.

## Introduction

Proper development and function of the immune system, digestive system, and metabolism require microbiota (Bosch and McFall-Ngai, 2021; Esser et al., 2019). Early-life exposure to microbial signals program the immune system and energy metabolism (Kimura et al., 2020; Torow et al., 2023). The immune system then reciprocally controls the composition of the microbiota by mucosal immunoglobulin A (IgA) and other regulatory mechanisms (Pabst et al., 2016). Interactions with microbes begin already before birth: maternal microbiota modulates fetal development by circulating metabolites (De Agüero et al., 2016; Husso et al., 2023) and possibly also by macromolecular components (Kaisanlahti et al., 2023). This prepares the immune system for the rapid microbial colonization at birth and for the transition towards adult diet and microbiota at weaning. Disturbance of the early host-microbe interactions can have long-lasting consequences on microbiota composition and host physiology (Schokker et al., 2015).

Maternally administered antibiotics can also perturb the microbiota in the offspring and increase the risk of immunological and metabolic disorders (Dierikx et al., 2020; Miyoshi and Hisamatsu, 2025). The dysbiotic microbiota of antibiotic-treated mothers at peripartum is transferred to the offspring (Miyoshi and Hisamatsu, 2025). However, little is known about the effects of prenatal maternal microbiota perturbation on fetal development. Experimental evidence is available mostly from mice, in which the short gestation does not allow the recovery from antibiotics before parturition, and therefore prenatal and postnatal effects cannot be distinguished. Complete lack of maternal microbiota during gestation caused broad changes in fetal gene expression and in postnatal intestinal innate immunity (De Agüero et al., 2016; Husso et al., 2023). In pigs, maternal antibiotic administration affected intestinal mucosa in the offspring (de Greeff et al., 2020), but in this study, the exposure occurred shortly before birth. In a prospective cohort study, 2^nd^ pregnancy trimester antibiotic exposure was associated with altered long-term microbiota composition and obesity in humans (Zhang et al., 2019).

Pigs serve as a valuable model for investigating human gut microbiota and immune system due to the physiological and microbial resemblance to humans (Heinritz et al., 2013; Pabst, 2020), while understanding the effects of maternal antibiotic treatment on piglet microbiota is important for animal husbandry. Use of antibiotics in pig farming has raised concerns about their effects on gut microbiota, antimicrobial resistance, and the overall health of the animals (Holman and Chenier, 2015). Investigating how maternal antibiotic treatment affects the microbiota of piglets is crucial, as the early colonization of microbes plays a significant role in immune system development and long-term health (Mueller et al., 2015; Schokker et al., 2015). Prior research has examined the impact of antibiotic exposure during pregnancy and early life on pigs, pointing out potential disruptions in microbial diversity and function (Guevarra et al., 2019). Antibiotic administration to sows during late pregnancy affected the development of offspring’s intestine and this change persisted for at least five weeks after birth (de Greeff et al., 2020). Weaning is a critical period for the development and growth of piglets and is accompanied by a large change in the microbiota, and alteration of the maternal microbiota can exacerbate dysbiosis at this stage (Guevarra et al., 2019; Madany et al., 2022b).

In this study, we exposed pregnant sows to narrow-spectrum (penicillin) or broad-spectrum (tetracycline) antibiotics at days 78-80 of the 114 to 119-day gestation. The aim was to perturb the maternal microbiota at a critical period in the fetal development of the pig immune system (Butler et al., 2017; Sinkora and Butler, 2009), allowing recovery of the material microbiota to pre-treatment status before parturition, and to investigate the effects of maternal antibiotic treatment on development of the intestinal microbiota and growth of the offspring.

## Methods

### Experimental design

This study was conducted on a commercial farm in Finland. The sows (Yorkshire × Landrace; n=98) had an average parity of 4.5 ± 1.6 (first parity sows were excluded) and had not received any antibiotics during their ongoing pregnancy. They were housed in two pregnancy rooms, both with partially slatted concrete floors. In room one, sows were kept in groups of 6 to 16, each with an individual feeding cage, while in room two, the group size ranged from 4 to 7, and they fed from long troughs. In both rooms, they had access to some roughage from a hay rack and water ad libitum from water nipples. At day 77 of gestation, they were weighed with a cage scale (DV203E digital, Danvaegt, Hinnerup, Denmark) and a veterinarian examined them clinically to include only healthy individuals not needing any medical treatments into the study. Body conditions were scored as (Norring et al., 2019): 1 = emaciated, 2 = thin, 3 = fit, 4 = fat and 5 = very fat. The sows were assigned systematically to three groups ensuring same parity distribution to each group. The following day, a three-day antibiotic treatment regimen was initiated, with sows receiving either an intramuscular injection of a saline solution (group CON, *Natriumchloride Braun Ecoflac* 9 mg/ml, 0.6 ml/10 kg) as a control, procaine penicillin (group PEN, *Ethacillin®* 300 mg/ml, 15 mg/kg), or tetracycline (group TET, *Engemycin LA®* 100 mg/ml, 5 mg/kg).

On day 113 of gestation, a second clinical examination was performed, and only healthy sows that had not received antimicrobial treatments after the first sampling were retained in the study. Out of 98 initially enrolled sows, 37 remained in the three study groups (CON n = 14, PEN n = 12, TET n = 11). To minimize the workload in farrowing supervision, 30 sows with farrowing during the same calendar days, were selected for further monitoring (10 per group; average parities: CON 4.4 ± 1.8, PEN 4.6 ± 1.8, TET 4.8 ± 1.8). The farrowings of the sows were supervised, and the birth time of the first piglet was recorded.

A total of 82 piglets [CON n = 30 (16 female, 13 male), PEN n = 24 (12 female, 12 male), TET n = 28 (14 female, 14 male)] from 19 sows (CON n = 7, PEN n = 6, TET n = 6) were included in the laboratory analyses, ensuring a balanced sex distribution across the sow medication groups. We only included piglets which received no antimicrobial treatments during the follow-up period and remained with their own dam. No piglets from other litters were moved into the study litters during the suckling period. After weaning, the study piglets were housed together in pens according to their dam’s medication group.

Piglets were weighed at one and three days as well as at three weeks of age using a digital hook scale (Pesola PHS040, China) and a custom-made fabric bag. At the age of ten weeks, they were weighed with the same cage scale, which was used for the sows earlier. Using weighting results we calculated average daily gain (ADG) for two growth periods: from birth to weaning (ADGsuckling) and from weaning to end of the follow-up period (ADGnursing) and these were used as outcomes to study microbiota associations with piglets weight gain.

Metadata for sows and piglets is included as Table S1. The animal experiment permit was granted by the Regional State Administrative Agency for Southern Finland (ESAVI/22394/2022).

### Sow sampling

Sows were sampled at two timepoints: before antibiotic administration at an average of 77 ± 0.2 days of gestation and again shortly before farrowing at 113 ± 0.2 days of gestation. Fecal and vaginal samples were collected at both stages, while colostrum samples were obtained within six hours of the birth of the first piglet. Vaginal samples were collected using sterile cotton swabs (Applimed SA, Châtel-St-Denis) and inserted into sterile milk sample tubes (Greiner Bio-One Gmbh, Germany). Fecal samples were obtained manually from rectum using factory clean gloves and placed in sterile 76 × 20 mm feces tubes (Sarstedt, Nümbrecht, Germany). Colostrum samples were collected following the procedure described in (Kaiser, 2019), after carefully wiping the nipple with disinfectant, into milk sample tubes (Greiner Bio-One Gmbh, Germany) and placed immediately after sampling in refrigerator temperature. After that the samples were aliquoted to sterile microcentrifuge tubes, immediately frozen to −20**°**C, and transferred to −80°C within one week.

### Piglet sampling

Piglets were sampled at three timepoints: at three days of age, just before weaning (three weeks = 23–27 days of age), and before the piglets were transferred to the finishing herd (ten weeks = 70–74 days of age). Fecal samples were collected using sterile cotton swabs (Applimed SA, Châtel-St-Denis) inserted into the rectum and placed into sterile cryotubes. All samples were placed immediately after sampling to refrigerator temperature, transferred to −20°C within one hour and transported in three days to laboratory storage in −80°C until analysis.

One of the day three samples in the CON group was excluded because the piglet was castrated before sampling. Four week ten samples in TET group and two in the PEN group were excluded due to antibiotic treatments before the last sampling. Thus, the final microbiota sample numbers were: day three, CON n = 29, PEN n = 24, TET n = 28; week three, CON n = 30, PEN n = 24, TET n = 28; week ten, CON n = 30, PEN n = 22, TET n = 24.

### DNA extraction and 16S rRNA gene amplicon sequencing

DNA was extracted from 125 mg of piglet feces, sow sampling swabs, and 200 μl of colostrum using the ZymoBIOMICS DNA Miniprep Kit based on mechanical cell lysis (Zymo Research, Irvine, CA), with minor modifications to the manufacturer’s protocol as described previously (Husso et al., 2020). Negative controls (DNA extracted from unused sampling instruments, and no-template controls), a ZymoBIOMICS Microbial Community Standard (Zymo Research), and an in-house complex microbiota standard (adult cow fecal DNA extract) were included. DNA concentration and purity was measured using a Nanodrop spectrophotometer 2000 (Thermo Fisher Scientific, Waltham, MA, USA).

For sequencing, the hypervariable regions V3 and V4 of the bacterial 16S rRNA genes were preamplified as described previously (Piirainen et al., 2025). Fecal and rectal swab samples were preamplified with 14 cycles, vaginal swabs with 18 cycles, colostrum samples with 21 cycles, along with negative controls for each sample type, and ZymoBIOMICS standard with 12 cycles. The sequencing was performed at the DNA Sequencing and Genomics Laboratory of the University of Helsinki, Finland. Excess primers were removed, and the indexing PCR was performed with 18 cycles. The PCR products were pooled and purified using MagSI-NGS plus beads (Magtivio, HK Nuth, Netherlands). The library pool was sequenced on the AVITI instrument (Element Biosciences, San Diego, CA) with paired-end reads (326 bp read 1 and 278 bp read 2) at a final library concentration of 7.5 pM, using the AVITI 2×300 Cloudbreak FS High Output Sequencing Kit.

### Amplicon sequencing data processing

Primers and spacers were trimmed using Cutadapt version 1.10 (Martin, 2011). The mapping file was validated using Keemei (Rideout et al., 2016). QIIME2 version 2024.2 (Bolyen et al., 2019) was used to process the sequencing data on the Puhti supercomputer of the CSC – IT Center for Science, Finland. Read quality was inspected and forward reads truncated to 243 bp and reverse reads to 204 bp to optimize data quality. The DADA2 (Callahan et al., 2016) plugin was used to merge and denoise the reads to amplicon sequence variants (ASVs). Taxonomic classification was performed using Greengenes2 version 2022.10 (McDonald et al., 2024) and a Naïve Bayes classifier trained on the target region. ASVs with less than 10 counts, mitochondrial and chloroplasts reads, and reads with undefined phylum were removed. The colostrum sequencing data was decontaminated using an R script, accepting ASVs which had at least 2× prevalence or at least 10× relative abundance in samples versus the negative controls. This removed 6.5% ± 5% of reads (mean ± standard deviation) in colostrum samples. The other sample types were not affected by contaminants due to their high microbial DNA content. The expected microbiota composition was observed for the ZymoBIOMICS Microbial Community Standard. The ASV count table is available as Table S2.

### Statistics

The microbiota compositions were analyzed in R version 4.5.0, using phyloseq 1.52.0 (McMurdie and Holmes, 2013), vegan 2.7-1 (Martinez Arbizu, 2020) and microbiome 1.30.0 (Lahti, 2017). Compositions were visualized using microViz 0.12.7 (Barnett et al., 2021). Alpha diversities were analyzed by the Shannon index in phyloseq, using non-rarefied ASVs with prevalence ≥5% in a sample type. Beta diversities were analyzed as Bray-Curtis distances of the relative abundances of species which were detected >10 times in ≥5% of samples and visualized using the plot_ordination function of phyloseq. PERMANOVA was performed using adonis2 and beta_disper of vegan, including group, sow parity, sow body condition score (BCS) (as a 4-level categorical variable), pregnancy room, farrowing room, and piglet sex.

Differential abundances were analyzed at genus level, in each sample type and each timepoint separately, using MaAsLin3 version 1.0.0, with total sum scaling (TSS, relative abundances) log_2_ transformation and Benjamini-Hochberg correction, without median comparisons (Nickols et al., 2024). Read depths and covariates which were significant in beta diversity analysis were included as fixed factors: group and sow parity (as three levels) in all analyses; for sow feces, also sow BCS (as four levels) and pregnancy room; for day three and week three piglets, also sow BCS, pregnancy room and farrowing room; and for week ten piglets, also sow BCS and piglet sex.

Longitudinal changes in genus abundances were analyzed by linear mixed-effects models (LMER) using the lme4 R package (Bates et al., 2015) for log-transformed relative abundances of genera with ≥25% prevalence and ≥0.1% mean relative abundance: lmer(abundance ∼ group * timepoint + parity + pregnancy_room + bcs + (1 | animal_ID), data = df, REML = TRUE, control = lmerControl (optimizer = “bobyqa”, optCtrl = list (maxfun = 2e5))). Type III ANOVA on primary effects and interactions was performed using lmerTest (Kuznetsova et al., 2017). P values were adjusted using the Benjamini–Hochberg correction (Benjamini and Hochberg, 1995).

Longitudinal changes in overall genus-level compositions were assessed as Aitchison distances of clr transformed data, using the volatility R package (Bastiaanssen et al., 2021). Genera which were detected >10 times in ≥5% of samples were included. Kruskal-Wallis and Wilcoxon tests were used for determining the significances of the observed differences.

The effects of treatment on alpha diversities and phylum-level compositions [log-transformed ratios of (Firmicutes+Actinobacteriota) / (Bacteroidota+Proteobacteria)] were analyzed by Generalized Linear Mixed Models package (GLMM) in SPSS 29.0.2.0. Variances explained by dam and pregnancy room were counted for by testing them as random intercepts to find the strategy resulting in the best model fit. Timepoint was included as repeated factor. Timepoint and treatment group as well as their interaction were included as fixed factors. Piglet sex as well as dam parity were controlled for if showing at least a statistical trend. Models were approved when homoscedasticity and normality of residuals could be established. Post-hoc pair-wise contrasts were interpreted with LSD adjustment for multiple comparisons.

Models were approved when homoscedasticity and normality of residuals could be established. Post-hoc pair-wise contrasts were interpreted with LSD adjustment for multiple comparisons.

General linear multivariate models (GLM) were used to analyze the associations between piglets’ fecal microbiota and ADG. Initially, to identify potential associations between sow-and piglet-related variables and piglet growth (analyzed separately for ADG during the suckling and nursing periods), a mixed linear regression model was applied, with sow included as a random intercept. After incorporating sow-related variables—such as parity, body condition score (BCS) group, and pregnancy room—as fixed factors, the random effect of the sow was no longer significant. Therefore, in subsequent models analyzing growth and ADG associations, these sow-related variables were included as fixed effects in all GLM models to control for potential confounding effects.

Four separate GLM models were constructed to investigate associations between fecal microbiota and piglet growth. For the suckling period, ADGsuckling was used as the outcome variable, with selected fecal genera at birth and at weaning as explanatory variables (two separate models). For the nursing period, ADGnursing was used as the outcome variable, with selected fecal genera at weaning and at the end of the study as explanatory variables (two additional models).

Potential confounders: sow parity (2–8), sow body condition score (BCS) group (<2.5, 3, 3.5, and 4), and pregnancy room (a two-level categorical variable) were included in all models.

To select the most relevant genera for inclusion in the multivariable GLM models, stability selection (Meinshausen and Bühlmann, 2010) was applied using the “stabiliser” package in R. Only genera with at least 10% prevalence and a minimum relative abundance of 0.1% were considered (at birth: n = 52; at weaning: n = 143; at the end of the study: n = 123). Sow parity, BCS, and pregnancy room were included as dummy variables.

A triangulation procedure (Lima et al., 2021), involving three variable selection models, was used to identify the most stable genera associated with ADG. The ten most stable genera were included in the final multivariate GLM models. Multicollinearity among explanatory variables was assessed. The linearity of the relationship between the outcome and continuous explanatory variables (genera relative abundances) was visualized using LOESS scatter plots. If a nonlinear relationship was observed, a log₂ transformation of genera relative abundances was applied. A stepwise backward elimination procedure was used for model selection, with a significance threshold of p ≤ 0.05. Model assumptions were evaluated by assessing residual normality and scatter plots.

To assess the potential confounding effect of sow antibiotic treatment on the associations between bacterial genera and ADG, the sow treatment group (CON, PEN, TET) was included in the final models. Changes in the coefficients of the genera were calculated, and a change of ≥10% was considered indicative of a confounding effect, suggesting that some of the associations between genera and ADG could be attributed to the sow’s antibiotic treatment.

The described GLM analyses were performed using preselected taxa based on their associations with ADG, as identified by the stability selection models. Taxa associated with the treatment group but not selected by the stability models were not included in these analyses.

To further explore associations between microbiota and growth, we conducted additional GLM analyses to assess the relationship between the relative abundance of bacterial genera and ADGsuckling or ADGnursing. These analyses focused on genera that exhibited significantly different relative abundances or significant timepoint × treatment group interactions at either three or ten weeks of age but were not among the 10 most stable genera identified by the stability selection models and thus were not included in the initial GLM models. In these additional models, sow-related confounders (parity, BCS, and pregnancy room) were included, and each genus was analyzed individually. The sow treatment group was also included in the final models to assess its potential confounding effect on the associations between these genera and ADG.

Similarly, we evaluated the associations between the Shannon diversity index and the Firmicutes+Actinobacteriota to Bacteroidota+Proteobacteria ratio with ADGsuckling and ADGnursing using GLM models. These models also included the sow treatment group to assess its potential confounding influence on the observed associations.

All GLM models were constructed using Stata/IC 14.2 (StataCorp, College Station, TX, USA). Generative AI (Claude Sonnet 4.5, Anthropic; and ChatGPT R Code Helper 5, OpenAI; both San Francisco, USA) was used for R coding assistance. All code was reviewed and validated by authors.

## Results

### Microbiota of sow feces, vagina, and colostrum

In sow fecal microbiota, there were no significant phylum or genus-level differences between the treatment groups (CON, PEN, and TET) at either timepoint (Figure 1). No significant differences were detected in alpha diversities between the treatment groups, nor were they different by beta diversity analysis.

**Figure 1.**
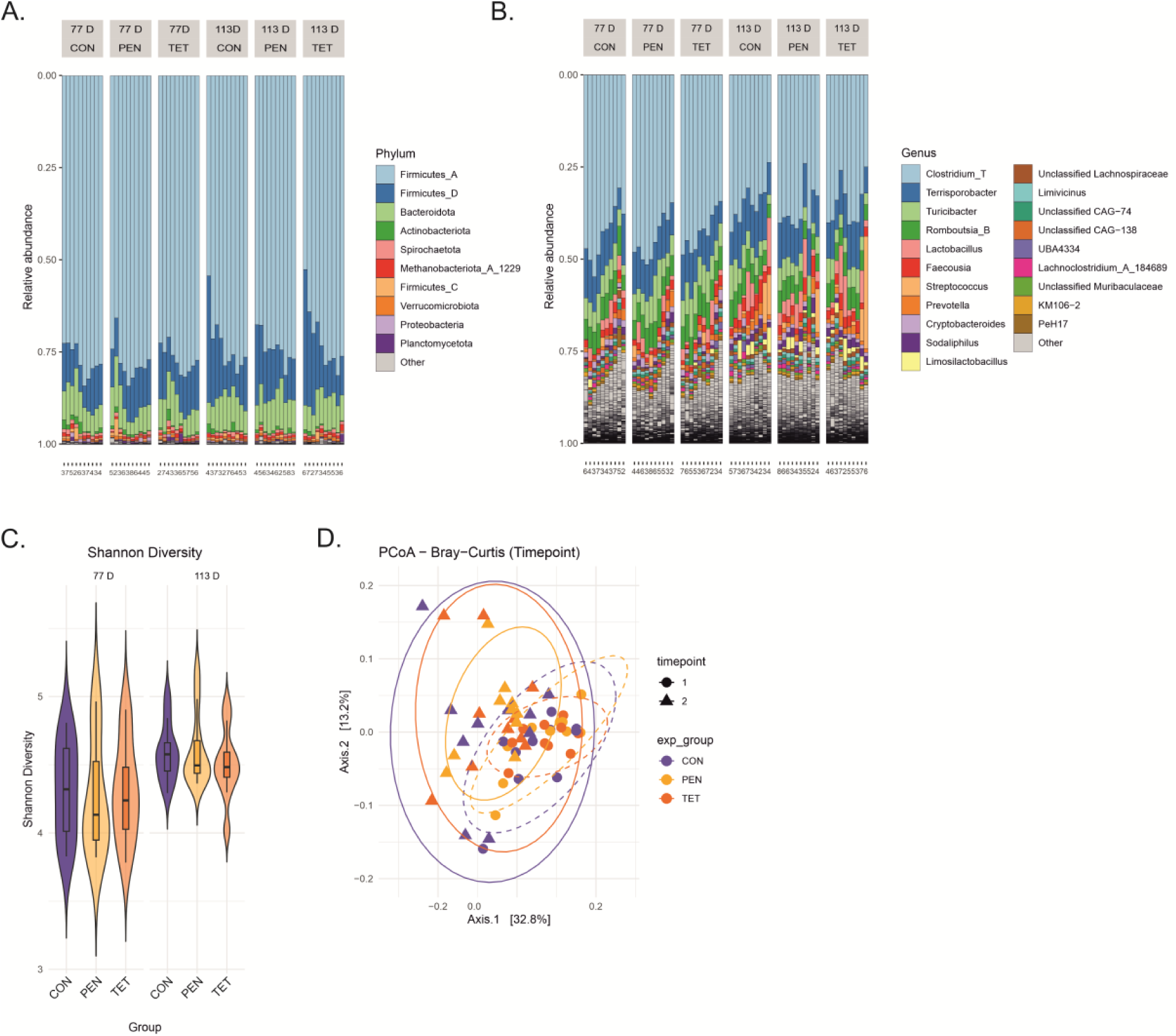
Fecal microbial profiles of pregnant sows treated with saline (CON = control group), penicillin (PEN) or tetracycline (TET) or during days 78-80 of pregnancy. Pregnancy day 77 (77 D) shows the results before treatments and 113 D shortly before farrowing. Microbiota compositions at (A) phylum and (B) genus level, numbers below the graph indicate parity. (C) alpha diversities of sow fecal microbiota (D) Beta diversity (PCoA plots of Bray-Curtis dissimilarity; 77D solid lines, 113D dashed lines).

A substantial change in sow fecal microbiota was however observed between the two phases of pregnancy. No significant differences in alpha diversity by Shannon were detected (Figure 1C) but the microbiota compositions were significantly different by beta diversity (ADONIS2 p < 0.004; Figure 1D). The relative abundances of 187 genera were significantly different between the sampling timepoints in LMER analysis, including lower abundances of the major genera *Clostridium_T*, *Romboutsia_B*, *Turicibacter*, and *Cryptobacteroides* and higher of *Sodaliphilus*, *Faecousia*, *Prevotella*, *Terrisporobacter,* and *Lactobacillus* at the end of pregnancy (Table S3).

Interestingly, in both antibiotic groups the fecal microbiota compositions changed significantly less between the sampling timepoints in comparison to CON, as measured by Aitchison distances; p = 0.043 for both comparisons.

In sow vaginal microbiota, the ratio of Firmicutes to Bacteroidota was significantly higher in TET vs CON at day 77 (Figure 2). No other significant differences were observed between the experimental groups at either phase of pregnancy by Shannon alpha diversity index, beta diversity, or genus-level abundances (Figure 2). The vaginal microbiota changed significantly between the phases of pregnancy by beta diversity (ADONIS2 p < 0.002; Figure 2D), but there was no significant difference in the magnitude of the change between the experimental groups. The relative abundances of *Fusobacterium_C* and *Veillonella_A* increased significantly from day 77 and both were among the most abundant at D 113, while the relative abundance of *Clostridium_AN*, one of the minor genera, decreased (Figure 2B and Table S4). Other significant changes were not detected.

**Figure 2.**
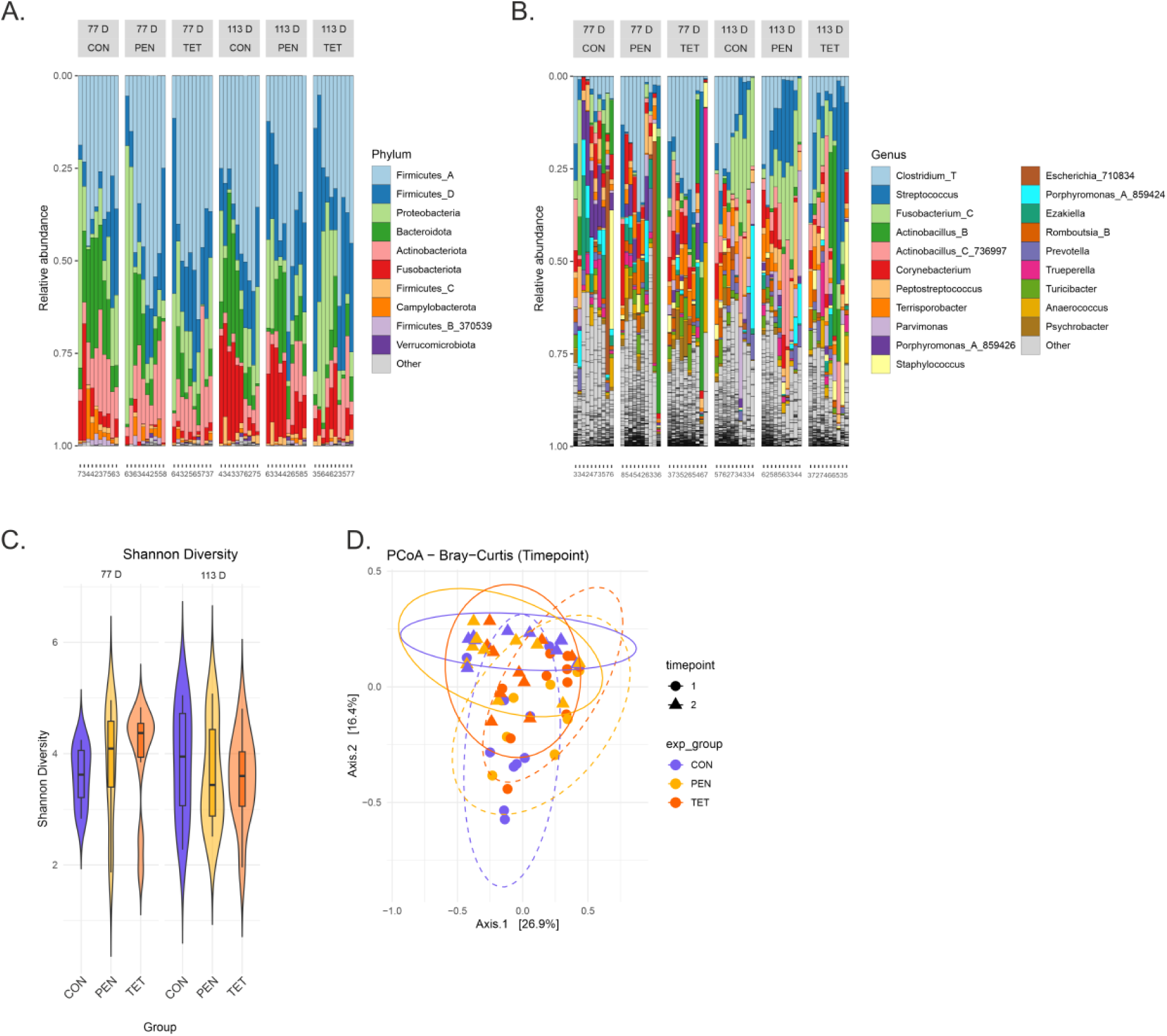
Vaginal microbial profiles of pregnant sows treated with saline (CON = control group), penicillin (PEN) or tetracycline (TET) during days 78-80 of pregnancy. Pregnancy day 77 (77 D) shows the results before treatments and 113 D shortly before farrowing. Microbiota compositions at (A) phylum and (B) genus level, numbers below the graph indicate parity. (C) alpha diversities of sow fecal microbiota (D) Beta diversity (PCoA plots of Bray-Curtis dissimilarity; 77D solid lines, 113D dashed lines).

In sow colostrum microbiota, neither the alpha diversity by Shannon nor beta diversity, nor the phylum ratio differed significantly between CON and the antibiotic-treated groups (Figure 3). No significant differences were detected in the microbial composition between the treatment groups at genus level. Three sows (two PEN and one TET sows) formed a separate cluster in the PCoA plot due to high abundances of *Streptococcus* as seen in Figure 6C (left). The ASVs matching the species *Streptococcus dysgalactiae* was very highly abundant in these three sows.

**Figure 3.**
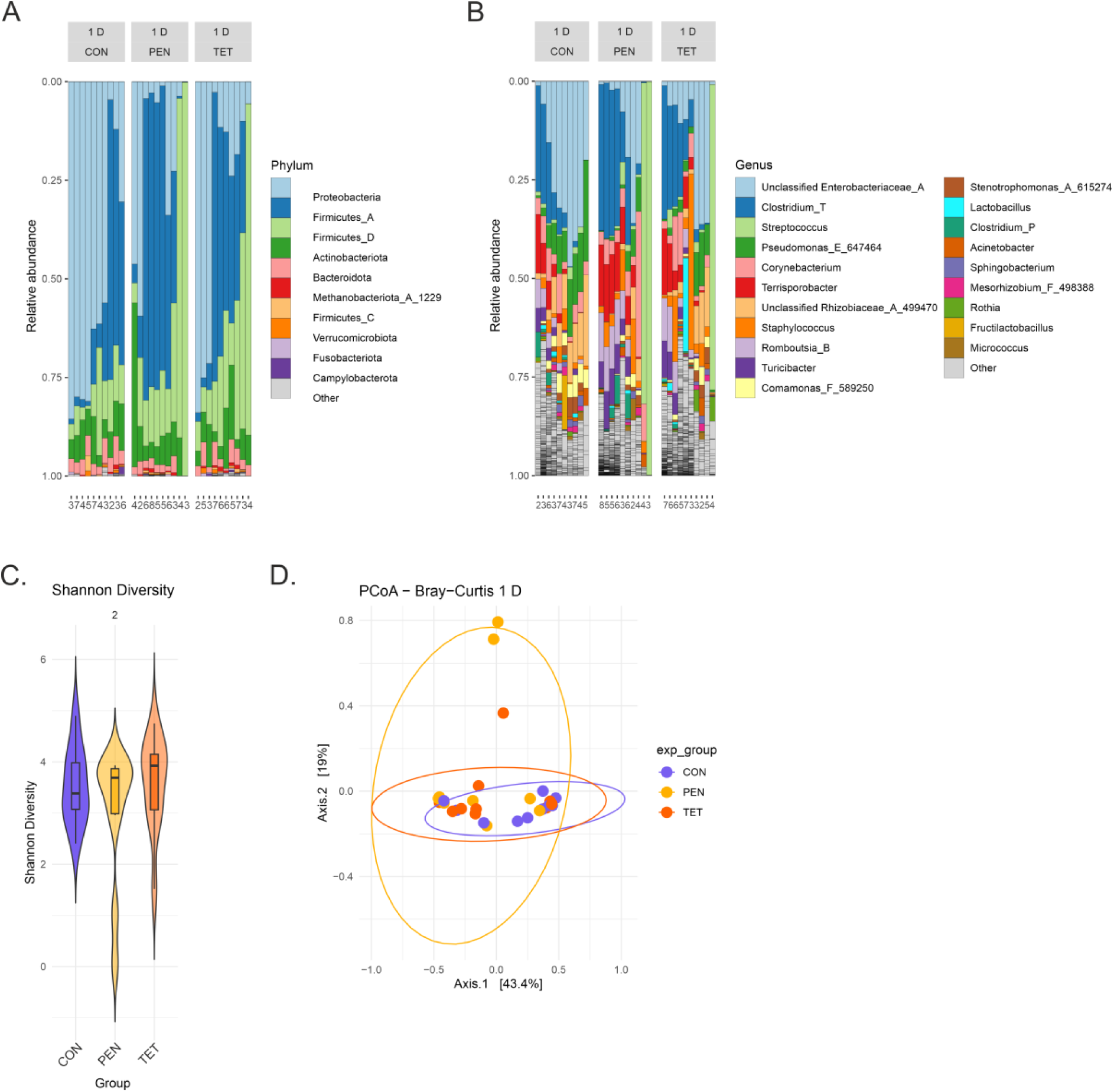
Colostrum microbial profiles within 6 hours after delivery of the first piglet in sows treated with saline (CON = control group), penicillin (PEN) or tetracycline (TET) during days 78-80 of pregnancy. Microbiota compositions at (A) phylum and (B) genus level, numbers below the graph indicate parity. (C) alpha diversities of sow fecal microbiota (D) Beta diversity PCoA plots of Bray-Curtis dissimilarity.

### Development of piglet fecal microbiota and impacts of maternal antibiotics

The microbiota of the 3-day-old piglets was characterized by high relative abundance of Proteobacteria (Figure 4A). *Clostridium_P, Escherichia_710834, Streptococcus,* and *Bacteroides_H* were the most abundant genera (Figure 4B). Alpha diversity was significantly lower in PEN versus CON (p = 0.007) (Figure 4C). The difference between CON and TET was not significant. The treatment groups differed in beta diversity (ADONIS2 p = 0.001). In PCoA of Bray-Curtis distances CON was slightly separated from PEN and TET (Figure 4D).

**Figure 4.**
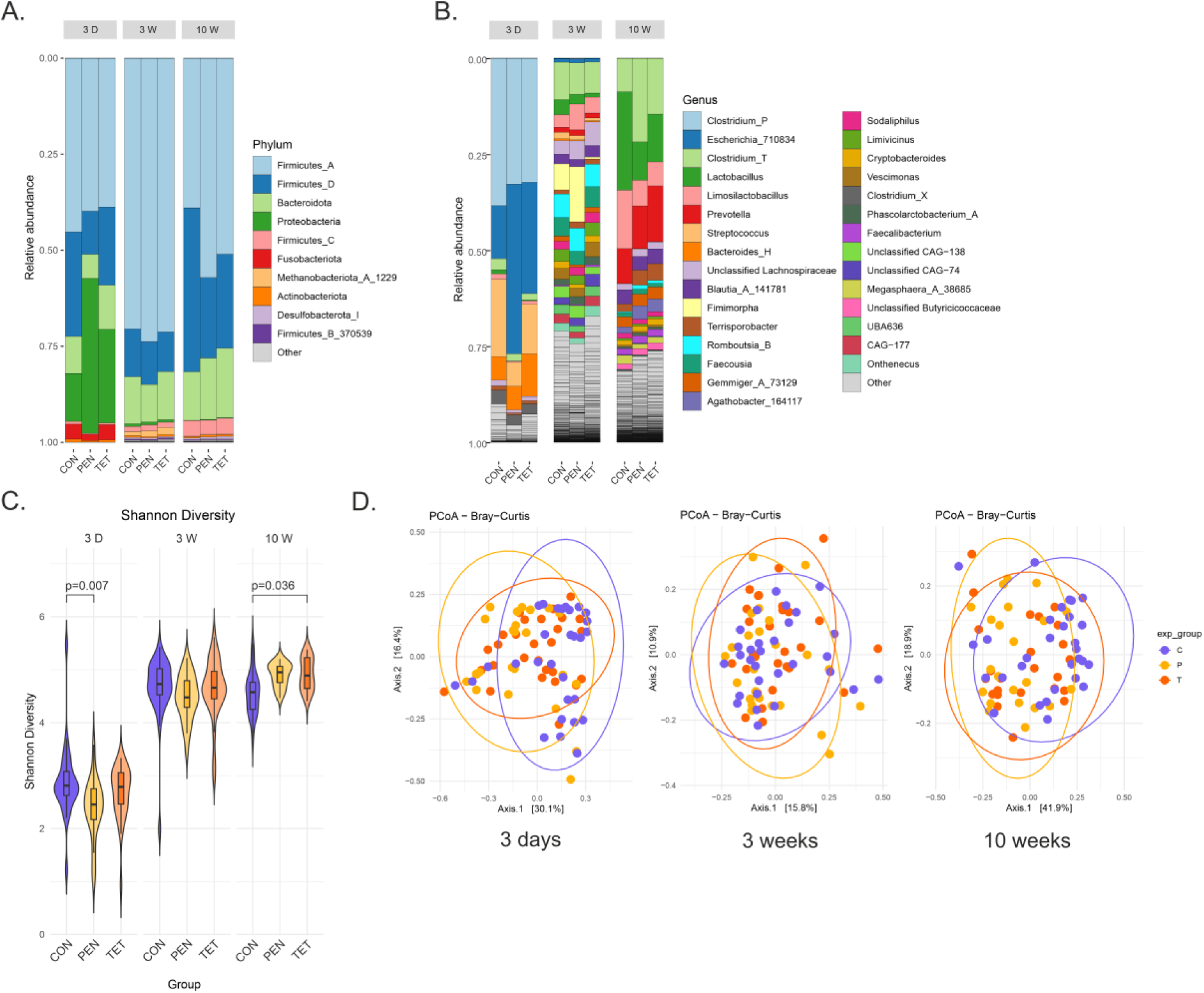
Piglet fecal microbiota composition and diversity at 3 days (3 D), 3 weeks (3 W) and 10 weeks (10 W) of age of **sows treated with saline (CON = control group), penicillin (PEN) or tetracycline (TET)during days 78-80 of pregnancy.** Microbial compositions at (A) phylum and (B) genus level. (C) Alpha diversity, as Shannon diversity index, (D) Beta diversity, as PCoA plots of Bray-Curtis dissimilarity.

The phylum ratio of (Firmicutes+Actinobacteriota) / (Bacteroidota+Proteobacteria) was significantly higher in CON vs PEN (p = 0.006) and CON vs TET (p = 0.016). At genus level, PEN had significantly higher abundance of *Escherichia _710834* and *Phocaeicola* _A_858004, and lower abundances of *Peptostreptococcus*, *Streptococcus*, Unclassified Cellulosilyticaceae, *Streptococcus*, and *Veillonella_A* in MaAsLin3 (Table S5). These genera also showed significant timepoint × treatment group interactions, i.e. differences in change between the treatment groups by time in LMER analysis (Figure 5 , Table S6). No significantly different genera were detected in TET vs CON.

**Figure 5.**
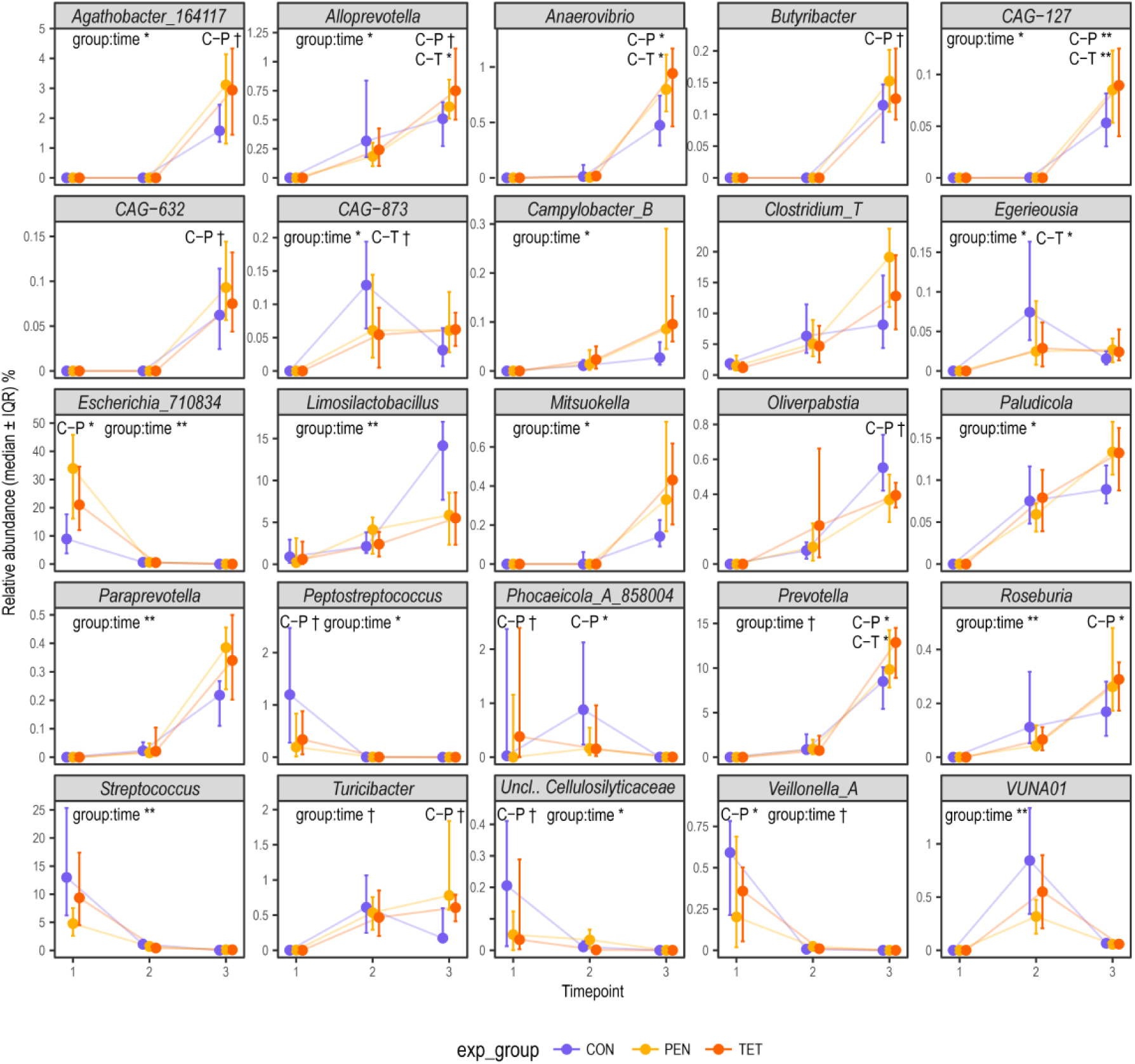
Trajectories of fecal microbial genera in piglets of sows treated with saline (CON = control group), penicillin (PEN) or tetracycline (TET) during days 78-80 of pregnancy. Horizontal axis age 3 days (3 D), 3 weeks (3 W) and 10 weeks (10 W), vertical axis relative abundance. Significance in MaAsLin3 at each timepoint and significance of timepoint × treatment group interaction in LMER are indicated in the plots: **\*\*** q < 0.01, *q < 0.05, † q < 0.1.

**Figure 6.**
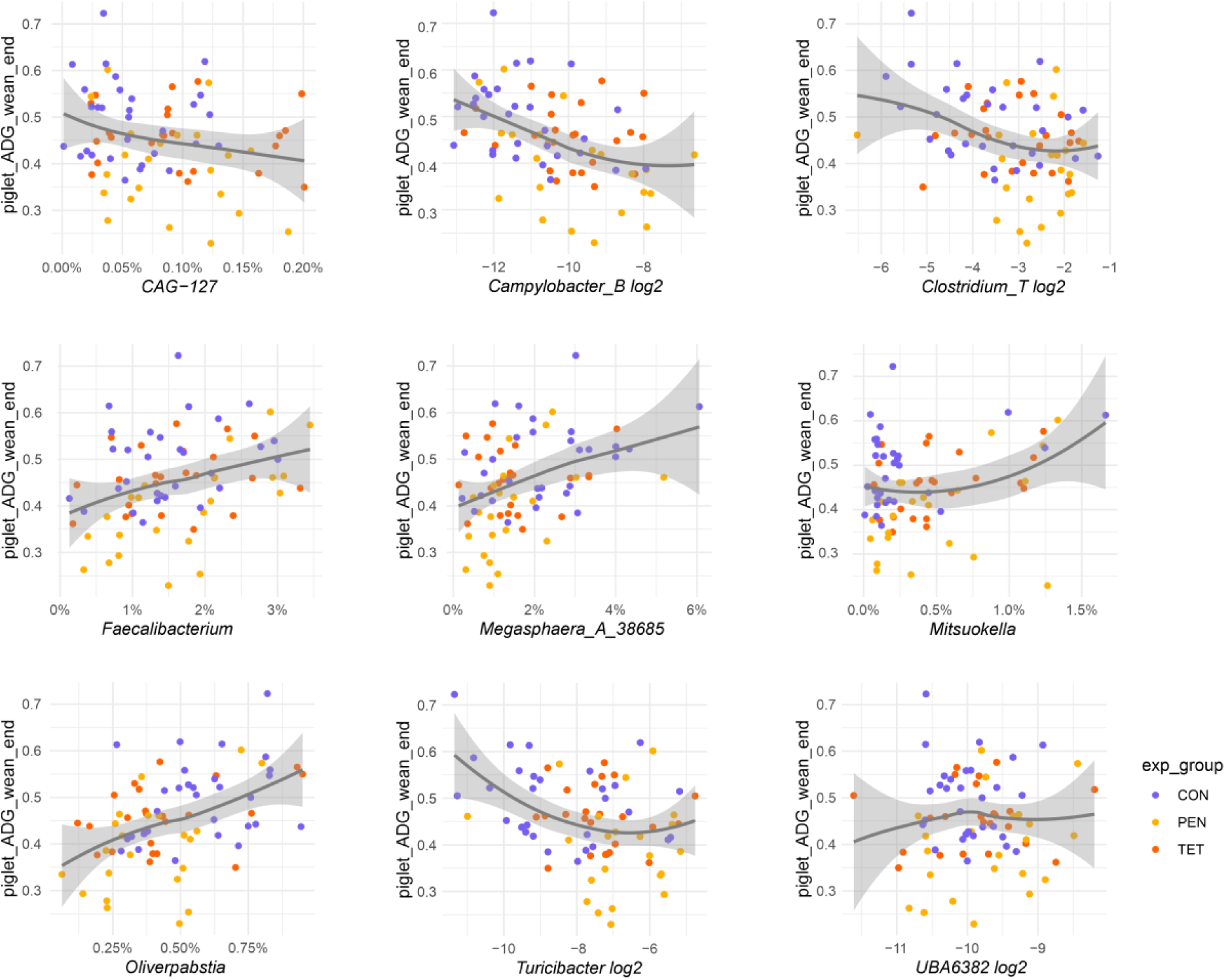
**Loess regression curves of association of relative abundances of bacterial genera at 10 weeks with ADGnursing in piglets of sows treated with saline (CON = control group), penicillin (PEN) or tetracycline (TET) during days 78-80 of pregnancy**.

In the end of the suckling phase, at three weeks of age, the most abundant phyla were Firmicutes_A, Firmicutes_D, and Bacteroidota, while Proteobacteria had much lower relative abundance (Figure 4A). *Clostridium_T, Fimimorpha, Romboutsia_B, Unclassified Lachnospiraceae, Limosilactobacillus* and *Faecousia* were the most abundant genera. (Figure 4B). Alpha diversity of the fecal microbiota of the piglets had increased during suckling (p 0.001. No significant differences in Shannon diversity were detected between the treatment groups (Figure 4C). The treatment group had significant effect by ADONIS2 (p = 0.008), but the groups overlapped in PCoA (Figure 4D).

The phylum ratio (Firmicutes+Actinobacteriota) / (Bacteroidota+Proteobacteria) tended to be lower in TET compared with CON (p = 0.097). At genus level, *Egerieousia* and *CAG-873* were significantly less abundant in TET vs CON, and *Phocaeicola_A_858004* more abundant in PEN vs CON by MaAsLin3 (Table S5). Significant differences were detected between the treatment groups in the change by time (timepoint × treatment group interactions) by LMER analysis for *Egerieousia* and *CAG-873,* and additionally for *Alloprevotella* and *VUNA01*, which were not significantly different at three weeks by MaAsLin3. (Figure 5 & Table S6).

At ten weeks the most abundant phyla were Firmicutes_A, Firmicutes-D, and Bacteroidota. Phylum Proteobacteria was almost absent (Figure 4A). *Lactobacillus*, *Clostridium_T*, *Prevotella*, *Limosilactobacillus, Blautia_A_141781, and Terrisporobacter* were the most abundant genera (Figure 4B). Alpha diversity by Shannon diversity index had increased further from week three to week ten (p = 0.017) and was higher in TET (p = 0.036) compared to CON (Figure 4C). No significant effects of treatment on beta diversity were detected by ADONIS2. CON separated partially from PEN in PCoA, TET overlapped with both the other groups (Figure 4D).

At ten weeks, the ratio Firmicutes + Actinobacteriota / Bacteroidota + Proteobacteria was higher in CON (p = 0.014) compared to TET (and tended to be higher in CON vs PEN (p = 0.063)). There were pronounced differences in the relative abundances of several genera between CON and the antibiotic groups. Both PEN and TET had significantly higher abundances of *Agathobacter_164117, Alloprevotella, Anaerovibrio, CAG-127,* and *Prevotella* than CON. Additionally, *Butyribacter*, *Roseburia*, and *Turicibacter* were significantly more abundant and *Oliverpabstia* less abundant in PEN vs CON in MaAsLin3 (Table S5). The abundance changes by time for *Agathobacter_164117, Alloprevotella, CAG-127, Roseburia*, and *Turicibacter* were significally different between the treatment groups (significant timepoint × treatment group interactions in LMER, Figure 5 & Table S6). In addition, the increase of the relative abundance of the major genus *Limosilactobacillus* from three to ten weeks was significantly stronger in CON than in the antibiotic groups (timepoint × treatment group interaction q = 0.003 in LMER, Figure 5 & Table S6).

The microbiota changed significantly more in TET compared to CON and PEN between three days and ten weeks (p.adj < 0.03 for Aitchison distances in both comparisons). In PEN, the change was not different from CON.

Piglet weight and growth

Piglets were born at an average weight of 1.48 ± 0.18 kg (SD), weaned at 8.51 ± 1.35 kg, and transferred to the fattening unit at 29.6 ± 5.1 kg. The ADGsuckling was 0.29 ± 0.05 kg/day (n = 80; birth weight was unavailable for two piglets), and ADGnursing 0.44 ± 0.10 kg/day (n = 82).

### Associations of piglet fecal microbiota with growth

We evaluated the effects of sow antibiotic treatment on the associations of piglet microbiota and growth. Treatment was included in all models evaluating these associations. There was a consistent and significant effect of sow antibiotic treatment on piglet ADGnursing across all GLM models evaluating associations between microbiota at three and ten weeks of age and weight gain. PEN piglets grew less than CON piglets, with model coefficients ranging from −0.129 to −0.074 kg/day and p-values between <0.001 and 0.031. TET group piglets also showed reduced growth compared to CON piglets, with coefficients ranging from −0.077 to −0.017 kg/day; however, p-values varied from 0.017 to 0.643 (see Tables S9 – S12). Sow antibiotic treatment was not associated with ADGsuckling in any of the models (see Tables S7 and S8).

We detected associations between microbial diversity and ADGnursing. An increase in Shannon diversity indices at three weeks by one unit was associated with an average increase of 0.037 kg/day in ADGnursing (SE = 0.017, p = 0.038), based on a GLM model that included sow-related variables (parity, BCS, pregnancy room) and piglet sex as confounders (n = 82). Adding treatment to this model did not substantially alter the coefficient (<10% change). Conversely, an increase in Shannon diversity at ten weeks (n = 76) by one unit was associated with an average decrease of 0.064 kg/day in ADGnursing (SE = 0.029, p = 0.029). Including treatment in this model decreased the coefficient by 17%, as Shannon diversities were higher in TET group than in CON (Figure 4) suggesting that the negative effect of Shannon diversity on weight gain was enhanced by sow antibiotic treatment (see Tables S13 and S14).

No associations were found between the phylum ratio of (Firmicutes + Actinobacteriota) / (Bacteroidota + Proteobacteria) at any of the three timepoints and ADGsuckling or ADGnursing (data not shown).

Associations of genus-level relative abundances at the three time-points to ADG were studied separately for the suckling and the nursery period using the GLM multivariable models (multiple genera were included as explanatory variables). Most notably, the relative abundances of *Oliverpabstia* and *Mitsuokella* at 10 weeks of age had significant positive associations with ADGnursing, and *CAG-127*, *UBA6382,* and *Campylobacter_B* had significant negative associations (Table S10, Figure 6). When treatment was included in the GLM model, the association coefficient for *Oliverpabstia* with ADG nursing decreased by 11%, while the negative association coefficient of CAG-127 with ADGnursing decreased by 33%, and the coefficient for *Campylobacter_B* decreased by 25% (Table S10). The relative abundance of *Oliverpabstia* was higher in the CON group compared to the PEN treatment group, while the relative abundance of *CAG-127* was higher in the PEN and TET groups compared to CON, and *Campylobacter_B* increased more in in the PEN and TET groups compared to CON (Figure 5).

Treatment also modulated the association coefficients of *UBA6382* and *Mitsuokella* with ADGnursing decreasing the coefficient of *UBA6382* by 36% and increasing that of *Mitsuokella* by 13%. However, their relative abundances were not significantly affected by the sow antibiotic treatment at 10 weeks by MaAsLin3 analysis (Figure 5).

The relative abundance of *Actinobacillus C 733574,* among a few other genera, at three days was negatively associated with ADGsuckling. Relative abundance of *Butyricimonas* at three weeks was negatively and *Romboutsia_B* positively associated with ADGsuckling. Relative abundances of *Paludicola* and *Sodaliphilus* at three weeks were positively associated with ADGnursing. However, inclusion of treatment to the GLM model had only small effects on the association coefficients of any of these genera (Tables S7-S9). *Faecousia* at three weeks was positively associated with ADGnursing, and inclusion of sow treatment in the model increased the association coefficient by 27%. In contrast, *Limousia* was negatively associated with ADGnursing, and adding the treatment variable decreased the coefficient by 19%. (Figure S1, Table S9).

Additional GLM models were constructed to evaluate genera that showed significantly different relative abundances between treatment groups or significant timepoint × treatment group interactions but were not among the 10 most stable genera identified by the stability selection models. These were included individually in the models to assess their associations with ADGnursing. Several of these genera showed significant associations, and sow antibiotic treatment was identified as a significant confounder in multiple cases (see Tables S11 and S12). LOESS regression curves illustrating these associations are presented in Figure 6.

The relative abundance of *VUNA01* at three weeks was significantly and positively associated with ADGnursing in the CON and PEN groups (p = 0.007) (Figure S1, Table S12). When treatment was included in the model, the association coefficient was reduced by 40%. Although the relative abundance of this genus was lower in the PEN and TET groups compared to CON, the difference was not statistically significant in MaAsLin3. However, a significant timepoint × treatment interaction was detected using LMER (Figure 5). *Alloprevotella* at three weeks showed a similar but weaker positive association with ADGnursing (p = 0.100). Including treatment in the model reduced the association coefficient by 30% (see Table S12).

*Turicibacter* had higher relative abundances in PEN than CON at ten weeks of age (MaAsLin3 q = 0.07). The genus had negative association with ADGnursing (p = 0.056), and the coefficient was increased by 33% when treatment was included. Similar results were obtained for *Clostridium_T*, which was one of the most abundant genera at ten weeks with higher relative abundance in PEN than CON (LMER q = 0.12 for the timepoint × treatment group interaction). This genus had significant negative association with ADGnursery (p = 0.018) and the coefficient increased by 15% when treatment was included (Table S11).

Two additional genera, *Faecalibacterium* and *Megasphaera_A_38685*, which have been shown to be positively associated with growth in earlier studies (Mahmud et al., 2023; Wang et al., 2023), were also included in the analysis. The relative abundance of *Faecalibacterium* at ten weeks had a positive association with ADGnursery (p = 0.041), which inclusion of treatment increased by 33% (Table S11). However, no effect on the relative abundance of this genus by antibiotic treatment was observed. The relative abundance of *Megasphaera_A_38685* at ten weeks also had a positive association with ADGnursery (p = 0.008), which was reduced 12% by inclusion of treatment (Table S11). At ten weeks it had lower abundances in PEN and TET vs CON, but the difference was not significant.

## Discussion

In this study, we treated pregnant sows with saline, penicillin, or tetracycline five weeks before parturition and analyzed their fecal, vaginal, and colostrum microbiota and the fecal microbiota of their offspring using 16S rRNA gene amplicon sequencing. No significant differences between the treatment groups were detected in sow fecal or vaginal microbiota compositions shortly before parturition at day 113 or in colostrum microbiota, but antibiotic treatment interfered with the dynamic change in maternal fecal microbiota during pregnancy. Significant differences in microbial diversities and compositions were observed in the fecal microbiotas of the piglets. The differences persisted and even increased after weaning and separation of the piglets from their dams, suggesting a possible impact on early programming of the mechanisms regulating host-microbe interactions. Microbial diversity at ten weeks was negatively associated with growth during the nursing period. Furthermore, relative abundances of several genera such as *Oliverpabstia*, *CAG-127* and *Campylobacter_B* at ten weeks and *Faecousia* and *Limousia* at three weeks were associated with growth. Antibiotic treatment modified these associations.

### Effect of antibiotic treatment on sow microbiota

Our results on the sow microbiota compositions show that the effects of the antibiotic treatment at day 77 of gestation on the fecal microbiota had subsided by day 113, as we expected. No major differences were observed between control and antibiotic-treated groups in fecal, vaginal or colostrum microbiota, making it unlikely that direct effects of antibiotics on sow microbiota compositions persisting until the time of delivery would have affected the microbiota of the piglets. While the microbial composition changed during pregnancy, we did not detect significant change in microbial richness, or in the relative abundances at phylum level between days 78 and 113. Substantial changes in the composition of the fecal microbiota are known to take place during pregnancy both in human and in pigs. The most robust differences involve decrease of the phyla Proteobacteria (Pseudomonadota) and Actinobacteria (Actinomycetota) in human (Koren et al., 2012) and significant reduction of microbial richness both in human and pigs (Kong et al., 2016; Koren et al., 2012) accompanied by differential abundances of several individual genera (Kong et al., 2016; Koren et al., 2012; Liu et al., 2025). The most significant differences are detected between early and late pregnancy. There was little overlap in the differentially abundant genera in our study compared to those reported in earlier studies with the exception of *Lactobacillus* (Kong et al., 2016; Liu et al., 2025) but, as shown by Liu et al. (2025), there can be considerable variation in the genus-level changes even in individual sows between pregnancies. However, the fecal microbiota changed less towards parturition in antibiotics-treated sows. Such changes may affect the early priming of the piglet immune system and the subsequent microbial colonization (Mueller et al., 2015; Schokker et al., 2015). The biological mechanisms of these interactions are still mostly unknown (Koren et al., 2024).

Previous studies on the effects of antibiotics during pregnancy have mainly focused on treatments given to sows at late pregnancy and/or during lactation (de Greeff et al., 2020; Ma et al., 2020; Wang et al., 2021; Wang et al., 2020; Zhu et al., 2024) directly modifying the sow intestinal and colostrum microbiota compositions during delivery and lactation (de Greeff et al., 2020). Further studies will be needed to determine how transient antibiotics-induced perturbations are mediated and affect the development of the piglet’s microbiota and its future health.

Vaginal microbiota is involved in the initial colonization of the newborn (Chen et al., 2022; Li et al., 2022; Liu et al., 2019) and changes in this microbiota may have implications for the piglet health. However, we did not detect any significant differences between the treatment groups that could be associated with the differences detected in the piglet fecal microbial compositions. Colostrum is another important source of microbes for colonization of the newborn piglet gut (Liu et al., 2019). We did not detect differences in the colostrum microbiota composition between treatment groups, indicating that antibiotic treatment about one month earlier had minimal impact on milk microbiota composition. A few sows belonging to the antibiotic groups showed high levels of *Streptococcus dysgalactiae*, a mastitis-associated bacterium, despite showing no clinical signs. These sows were not among those whose piglets were included in the study. Similar findings were made in sow colostrum in an earlier study not involving the use of antibiotics (Piirainen et al., 2025).

### Early-life microbiota: Initial colonization

Earlier research has shown that antibiotic treatment of dams during pregnancy has impacts on the development of the commensal microbiota of their offspring (de Greeff et al., 2020; Dierikx et al., 2020; Zhang et al., 2019). In most experimental animal studies addressing this question, antibiotics were administered orally shortly before or at the time of parturition (Alhasan et al., 2020; Alhasan et al., 2023; Champagne-Jorgensen et al., 2020; Chen et al., 2021; de Greeff et al., 2020; Gonzalez-Perez et al., 2016; Madany et al., 2022a, b; Wang et al., 2021; Zou et al., 2018) and often also during lactation (Alhasan et al., 2020; Chen et al., 2021; Wang et al., 2020). Effects often included reduced microbial richness and lower relative abundance of Firmicutes and Actinobacteria in relation to Proteobacteria in the early gut microbiota of the offspring (Chen et al., 2021; Gonzalez-Perez et al., 2016; Zou et al., 2018) Analogous findings have been recorded in many studies on the effect of maternal therapeutic antibiotics on the microbiota development of human infants (Dierikx et al., 2020). Interestingly, we observed similar effects on the developing microbiota of the offspring when the dams were treated with antibiotics intramuscularly as early as five weeks before parturition. At three days of age the fecal microbiota of piglets of PEN-treated sows had lower alpha diversity and lower relative abundance of the phyla Firmicutes + Actinobacteriota in relation to Bacteroidota + Proteobacteria.

Significant differences in bacterial abundance were detected between the treatment groups. The PEN group piglets had higher relative abundances of *Escherichia_710834* and *Phocaeicola*_*A_858004*. While there is considerable variation in taxa reported to be differentially abundant in the offspring without or with maternal antibiotic treatment, higher relative abundance of the genus *Escherichia* or family Escherichia-Shigella has been reported in the newborn progeny of antibiotic-treated vs control in mice and human (Alhasan et al., 2023; Chen et al., 2021; Zou et al., 2018). Species belonging to this taxon have been linked to dysbiosis and opportunistic infections (Denamur et al., 2021; Luo et al., 2022; Pantazi et al., 2025). The PEN group piglets also had lower abundances of *Peptostreptococcus*, *Streptococcus*, Unclassified Cellulosilyticaceae, and *Veillonella_A* than controls. *Peptostreptococcus* and *Veillonella* have been reported to belong to the most abundant genera in weaned piglets with diarrhea (Kong et al., 2022). The genus *Streptococcus* contains several potentially pathogenic species, such as *S. suis* and *S. dysgalactiae* (Ma et al., 2025) but the ASVs matching these species were not significantly different between the treatment groups.

### Pre-weaning transition: three weeks

At three weeks of age, the differences between the antibiotic groups and the controls had subsided. The alpha diversity had increased substantially in all the groups and no effects of antibiotic treatment on alpha diversity measures were detectable. There was only a tendency to lower phylum ratio in TET vs CON. The genera *Phocaeicola*, *CAG-873*, *VUNA01* and *Alloprevotella* were significantly differentially abundant between the groups. *Phocaeicola_A_858004*, which had lower abundance in the antibiotic groups, is a gut commensal identified as *P. vulgatus,* which is recognized as a potential carbohydrate utilizer in pigs (Holman et al., 2022). The species has been shown to utilize milk oligosaccharides *in vitro* (Rumeau et al., 2025). *Egerieousia,* with lower abundance in TET, is a relatively uncharacterized genus that was identified by metagenomic analysis in chicken gut (Gilroy et al., 2021). *Egerieousia* sp004561775 has been identified as a potential butyrate producer in pigs (Holman et al., 2022). Little information is available on *Prevotella CAG-873,* also less abundant in TET, except that a strong positive correlation of sow *Akkermansia* during lactation to piglet fecal *CAG-873* after weaning has been shown (Saladrigas-Garcia et al., 2022). *VUNA01* is a bacterial taxon identified in pigs (Holman et al., 2022) but its properties are not known.

### Post-weaning: maturation

By ten weeks the alpha diversity had increased further towards more mature levels in all groups but was significantly higher in TET compared to CON. The Firmicutes + Actinobacteriota / Bacteroidota + Proteobacteria ratio was lower in PEN than CON. The relative abundance of *Oliverpabstia* was significantly higher in CON than in the antibiotic groups. In addition, the increase in the relative abundance of *Limosilactobacillus* by ten weeks was much more pronounced in CON than in the antibiotic groups. Several genera, including *Agathobacter_164117, Alloprevotella, Anaerovibrio, Butyribacter, CAG-127, Roseburia*, *Prevotella, and Turicibacter* were more abundant in one or both antibiotic groups.

*Limosilactobacillus* is a genus containing potential probiotic species (Ksiezarek et al., 2022). Species belonging to this genus have been shown to improve the intestinal barrier function in piglets (Li et al., 2023) and other species (Wu et al., 2022). *Oliverpabstia* (Wylensek et al., 2020) is a genus of the family Lachnospiraceae that contains species enriched in piglet feces after weaning (Holman et al., 2022).

*Agathobacter, Anaerovibrio, Alloprevotella, Prevotella,* and *Turicibacter* are all relatively abundant commensal bacteria. *Anaerovibrio* contains species with lipolytic activity growing on sugars (Strömpl et al., 1999)*. Alloprevotella* utilizes carbohydrates and produces succinic acid (Downes et al., 2013)*. Agathobacter* (Rosero et al., 2016) and *Prevotella* (Ivarsson et al., 2014) are involved in fiber degradation and carbohydrate metabolism and production of short-chain fatty acids. The relative abundance of *Prevotella* has been reported to increase after weaning in piglets (Guevarra et al., 2018; Mach et al., 2015), with positive correlations of its relative abundance to body weight and soluble IgA concentration (Mach et al., 2015). Decreased abundance of *Prevotella* in the offspring after maternal antibiotic treatment has been reported in several studies on mice and human (Alhasan et al., 2023; Champagne-Jorgensen et al., 2020; Zhu et al., 2024). *Turicibacter* has the capability of decreasing the levels of lipids and modulating bile acid profiles of in the serum of mammalian hosts (Lynch et al., 2023)*. CAG-127* is a member of the family Lachnospiraceae known for production of short-chain fatty acids (Brito Rodrigues et al., 2025).

### Associations with growth performance

The associations of microbiota and growth were strongest after weaning. *Oliverpabstia* and *Mitsuokella* were positively associated with growth, and *CAG-127* and *Campylobacter_B* negatively associated. Inclusion of sow antibiotic treatment to the statistical models had substantial effects on the association coefficients. The results suggests that antibiotic treatment reduced the positive effect of *Oliverpabstia* and enhanced the negative effect of *CAG-127* and *Campylobacter_B* on ADG-nursing. As the relative abundance of *Oliverpabstia* was higher in the CON group compared to the PEN treatment group, the positive effect of this genus on growth in CON vs PEN may be due to its higher relative abundance in CON. The relative abundances of *CAG-127* and *Campylobacter_B* were higher in the PEN and TET groups compared to CON, and their negative effect on growth during the nursery period may be due to their higher abundances in the antibiotic groups. Although treatment also modulated the association coefficients of *UBA6382* and *Mitsuokella* with ADGnursing decreasing the coefficient of *UBA6382* and increasing that of *Mitsuokella,* their relative abundances in piglets were not significantly affected by the sow antibiotic treatment. Therefore, the effect of treatment on growth through these genera cannot be interpreted directly. However, the results suggest that sow antibiotic treatment may have enhanced the negative impact of *UBA6382* and increased the positive impact of *Mitsuokella* on growth during the nursery period.

*Turicibacter* and *Clostridium_T* at ten weeks had significant negative association to ADGnursery, and treatment increased the coefficient, suggesting that treatment enhanced their negative impact on growth. The relative abundances of these genera were higher in PEN than CON, suggesting a link between the abundance and the negative effect. In contrast, *VUNA01* at three weeks had a positive association with growth during the nursery period, which was reduced by antibiotic treatment. Similar observation was made for *Alloprevotella* though its association with growth was weaker.

The positive association of *Faecousia* at three weeks with ADGnursing was increased and the negative association of *Limousia* with ADGnursing was decreased by inclusion of sow treatment in the model. These results suggest that sow antibiotic treatment enhanced the positive effect of *Faecousia* and intensified the negative effect of *Limousia* on growth. However, their relative abundances did not differ significantly between the treatment groups.

To our knowledge, associations of *Oliverpabstia* or *CAG-127* with piglet growth, or maternal antimicrobial influence on their abundances in piglet gut have not been reported earlier. However, we have observed a significantly higher relative abundances of *Mitsuokella* in high ADG weaning piglets in an earlier study (Mahmud et al., 2023). Moreover, a positive association of *Mitsuokella* with body weight and ADG in piglets has been reported (Yang et al., 2021). *Campylobacter* is a genus containing intestinal pathogens and it has been associated with post-weaning diarrhea in piglets (Yang et al., 2020; Zheng et al., 2023) with accompanying metabolic dysfunction (Yang et al., 2020). *Oliverpabstia* and *Faecousia* have been implicated in maternal obesity in a rat model (Li et al., 2025). *Turicibacter* belongs to the genera reported to be related to post-weaning transition of piglets (Yi et al., 2025). The major ASV corresponding to *Clostridium_T* in our data was identified as *C. disporicum*, which is a saccharolytic commensal species (Horn, 1987) with pathogenic potential (Woo et al., 2005).

### Limitations of the study

The sows were given antibiotics at days 78-80 of gestation, and the subsequent sampling was done only on day 113. Hence, any short-term alterations in microbiota, that were likely to have taken place following the treatment, could not be traced. Secondly, we employed 16S rRNA gene amplicon sequencing to assign taxonomy to near-genus level, which is less discriminatory than shotgun metagenomic sequencing. Lastly, the experiment was carried out on a commercial farm, with many environmental and farm management factors inducing experimental variation. This may have prevented detection of some treatment effects and associations due to insufficient statistical power. Despite these limitations, this research provides valuable insights into the effects of maternal antibiotic exposure on neonatal gut microbiota and lays the groundwork for future studies.

## Conclusion

Our study highlights that maternal antibiotic use during pregnancy has a long-term impact on the gut microbiota of piglets and impairs growth, even if the maternal microbiota largely recovers by parturition. Significant treatment-related differences in the fecal microbial compositions were evident in the piglets, especially at ten weeks of age. This suggests that maternal antibiotic exposure affects not only the establishment of the initial gut microbiota but also its progressive development in the offspring.

Maternal antibiotic treatment in the beginning of the last third of pregnancy did not significantly alter the microbiota of sows at farrowing but impaired the dynamic development of the fecal microbiota over gestation, which may affect the early priming of the piglet immune system and subsequent microbial colonization. This study stresses the importance of cautious antibiotic use during pregnancy in livestock to protect offspring health and productivity. Future research should explore interventions such as probiotics or modulation to balance the microbiome in piglets.

## Acknowledgements

We thank Kirsi Lahti for expert technical assistance. We acknowledge DNA Sequencing and Genomics Laboratory (supported by HiLIFE and Biocenter Finland funding), Institute of Biotechnology, University of Helsinki for sequencing, and CSC – IT Center for Science, Finland, for generous computational resources.

## Disclosure statement

No potential conflict of interest was reported by the authors.

## Funding

This work was supported by the Finnish Ministry of Agriculture and Forestry (project number VN/7587/2021), Research Council of Finland (project 347925), A-Producers Ltd, and Vetcare Ltd. Open access is funded by Helsinki University Library.

## Author contributions

Conceptualization and design the study: M.H., M.N., K.A., S.J., T.P.M.; sample collection: M.R.M., M.H., K.A., E.K., S.J., M.N., T.P.M., H.N.K.; DNA extraction and quantification: M.R.M.; metadata preparation: M.H., K.A., M.N., M.R.M.; bioinformatics analysis, M.R.M & M.N.; statistical analysis: M.R.M., M.N., T.O, C.M., statistical analysis supervised M.P, M.L.C, M.N., T.O.; manuscript writing: M.R.M. T.P.M, M.H., T.O. C.M.; editing and reviewing manuscript: M.N, T.P.M, M.H., C.M., T.O, K.A., S.J., C.O., E.K. C.M; supervision, M.N., M.H., C.O. Approval of the final manuscript: All the authors read and approved the final manuscript.

## Data availability

The 16S rRNA gene amplicon sequencing data will be available in the European Nucleotide Archive (ENA) repository under accession no. PRJEB100560.

## Notes

### Competing Interest Statement

The authors have declared no competing interest.

### Summary of Updates

This version of the manuscript has been revised to update some missing references and correct a few typing errors.

