## Supplementary Tables S7-S14 for "Impact of gestational antibiotics on maternal and offspring gut microbiota and growth in pigs"

**Table S7.** Results from a multivariable general linear model evaluating associations between piglets' (n = 79) fecal microbiota relative abundace at the genus level at three days and average daily weight gain (kg/day) from birth to the weaning (ADGsuckling), both before and after including the sow treatment group to assess potential confounding effects.

| Variable | Coeff. | SE | p-value | Wald test p-value |
|---|---|---|---|---|
| **Sow parity group:**[1] 2-8 | | | | 0.384* |
| **Sow body condition score group:**[1] 2.5, 3, 3.5 and 4 | | | | 0.708* |
| **Sow pregnancy room:** | | | | |
| 1 | 0 | | | |
| 2 | 0.073 | 0.023 | 0.001* | |
| Actinobacillus C 733574 | -15.487 | 6.284 | 0.014 | |
| Clostridium P (log2) | -0.131 | 0.037 | <0.001 | |
| Actinobacillus B (log2) | -0.916 | 0.339 | 0.007 | |
| Intercept | 0.264 | 0.036 | <0.001 | |
| **ADG-total and genus associations after icluding sow treatement group** | | | | **Change of coeff.** |
| **Sow treatment group:** | | | | |
| No treatment | 0 | | | |
| Penicillin | -0.017 | 0.023 | 0.456 | |
| Tetracycline | -0.028 | 0.022 | 0.210 | |
| Actinobacillus C 733574 | -15.661 | 6.315 | 0.013 | 0.5% |
| Clostridium P (log2) | -0.130 | 0.037 | <0.001 | 1% |
| Actinobacillus B (log2) | -0.949 | 0.351 | 0.007 | 3% |
| Intercept | 0.235 | 0.045 | <0.001 | |

[1] Group differences are omitted from the table

**Table S8.** Results from a multivariable general linear model evaluating associations between piglets' (n = 80) fecal microbiota relative abundace at the genus level at weaning and average daily weight gain (kg/day) from birth to the weaning (ADG-suckling), both before and after including the sow treatment group to assess potential confounding effects.

| **Variable** | **Coeff.** | **SE** | **p-value** | **Wald test p-value** |
|---|---|---|---|---|
| **Sow parity group:**[1] 2-8 | | | | 0.362* |
| **Sow body condition score group:**[1] 2.5, 3, 3.5 and 4 | | | | 0.070* |
| **Sow pregnancy room:** | | | | |
| 1 | 0 | | | |
| 2 | -0.013 | 0.022 | 0.535* | |
| Romboutsia_B | 0.461 | 0.109 | <0.001 | |
| Butyricimonas | -7.249 | 2.795 | 0.009 | |
| Intercept | 0.291 | 0.038 | <0.001 | |
| **ADG-sucling and genus associations after icluding sow treatement group** | | | | **Change of coeff.** |
| Sow treatment group: | | | | |
| No treatment ( | 0 | | | |
| Penicillin | -0.032 | 0.022 | 0.148 | |
| Tetracycline | -0.041 | 0.022 | 0.061 | |
| Romboutsia B | 0.475 | 0.111 | <0.001 | 2% |
| Butyricimonas | -7.656 | 2.797 | 0.006 | 5% |
| Intercept | 0.245 | 0.046 | <0.001 | |

[1] Group differences are omitted from the table

* Variable is included to control the confounding effect

**Table S9.** Results from a multivariable general linear model evaluating associations between piglets' (n = 82) fecal microbiota relative abundace at the genus level at weaning and average daily weight gain (kg/day) from weaning to the end of the study period (ADG-nursing), both before and after including the sow treatment group to assess potential confounding effects.

| **Variable** | **Coeff.** | **SE** | **p-value** | **Wald test p-value** |
|---|---|---|---|---|
| **Sow parity group:**[1] 2-8 | | | | <0.001* |
| **Sow body condition score group:**[1] 2.5, 3, 3.5 and 4 | | | | <0.001* |
| **Sow pregnancy room:** | | | | |
| 1 | 0 | | | |
| 2 | -0.02 | 0.031 | 0.509* | |
| Sodaliphilus | 0.977 | 0.399 | 0.015 | |
| Faecousia | 0.673 | 0.295 | 0.023 | |
| Limousia | -1.287 | 0.297 | <0.001 | |
| Paludicola | 44.437 | 18.876 | 0.019 | |
| Intercept | 0.598 | 0.056 | <0.001 | |
| **ADG-nursing and genus associations after icluding sow treatement group** | | | | **Change of coeff.** |
| **Sow treatment group:** | | | | |
| No treatment | 0 | | | |
| Penicillin | -0.074 | 0.034 | 0.031 | |
| Tetracycline | -0.046 | 0.032 | 0.152 | |
| Sodaliphilus | 0.891 | 0.393 | 0.023 | 8% |
| Faecousia | 0.493 | 0.304 | 0.105 | 27% |
| Limousia | -1.043 | 0.312 | 0.001 | 19% |
| Paludicola | 45.634 | 18.504 | 0.014 | 3% |
| Intercept | 0.533 | 0.067 | 0 | |

[1] Group differeces are omitted from the table

* Variable is included to control the confounding effect

**Table S10.** Results from a multivariable general linear model evaluating associations between piglets' (n = 76) fecal microbiota relative abundace at the genus level at the end of the study period and average daily weight gain (kg/day) from weaning to the end of the study period (ADG-nursing), both before and after including the sows treatment group to assess potential confounding effects.

| Variable | Coeff. | SE | p-value | Wald test p-value |
|---|---|---|---|---|
| **Sow parity group:**[1] 2-8 | | | | <0.001* |
| **Sow body condition score group:**[1] 2.5, 3, 3.5 and 4 | | | | <0.001* |
| **Sow pregnancy room:** | | | | |
| 1 | 0 | | | |
| 2 | 0.008 | 0.038 | 0.820* | |
| Oliverpabstia | 8.775 | 4.278 | 0.040 | |
| CAG-127 | -36.599 | 16.931 | 0.031 | |
| Campylobacter B ($log_2$) | -0.025 | 0.012 | 0.039 | |
| Mitsuokella | 6.177 | 2.355 | 0.009 | |
| UBA6382 ($log_2$) | -0.062 | 0.026 | 0.019 | |
| Intercept | -0.219 | 0.299 | 0.464 | |
| **ADG-nursing and genus associations after including sow treatement group** | | | | **Change of coeff.** |
| **Sow treatment group:** | | | | |
| No treatment | 0 | | | |
| Penicillin | -0.091 | 0.038 | 0.015 | |
| Tetracycline | -0.017 | 0.037 | 0.643 | |
| Oliverpabstia | 7.771 | 4.048 | 0.049 | 11% |
| CAG-127 | -24.953 | 18.226 | 0.171 | 32% |
| Campylobacter B ($log_2$) | -0.017 | 0.011 | 0.135 | 32% |
| Mitsuokella | 7.429 | 2.223 | 0.001 | 17% |
| UBA6382 ($log_2$) | -0.038 | 0.026 | 0.136 | 39% |
| Intercept | 0.035 | 0.292 | 0.902 | |

[1] Group differeces are omitted from the table

* Variable is included to control the confounding effect

**Table S11.** Results from univariable general linear models evaluating associations between piglets' (n = 76) fecal microbiota relative abundace at the genus level at the end of the study period and average daily weight gain (kg/day) from weaning to the end of the study period (ADG-nursing), both before and after adjusting for the sows' treatment group to assess potential confounding effects. All potential confounding variables (sow parity, sow body contition score and pregnacy room) were included in the univariable models (results not shown). The genera selected for the univariable models did not show significant associations with weight gain in the multivariable model, but were significantly associated with the sows' treatment group.

| Variable | Coeff. | SE | p-value | Change of coeff. |
|---|---|---|---|---|
| **Turicibacter ($log_2$)** | -0.013 | 0.008 | 0.100 | |
| **Sow treatment group:** | | | | |
| No treatment | 0 | | | |
| Penicillin | -0.125 | 0.035 | <0.001 | |
| Tetracycline | -0.059 | 0.035 | 0.090 | |
| **Turicibacter ($log_2$)** | -0.008 | 0.007 | 0.274 | 33% |
| **Clostridium T ($log_2$)** | -0.021 | 0.009 | 0.019 | |
| **Sow treatment group:** | | | | |
| No treatment | 0 | | | |
| Penicillin | -0.126 | 0.034 | <0.001 | |
| Tetracycline | -0.063 | 0.034 | 0.063 | |
| **Clostridium T ($log_2$)** | -0.018 | 0.008 | 0.031 | 15% |
| **Megasphaera A 38685** | 2.090 | 0.785 | 0.008 | |
| **Sow treatment group:** | | | | |
| No treatment | | | | |
| Penicillin | -0.122 | 0.033 | <0.001 | |
| Tetracycline | -0.054 | 0.033 | 0.109 | |
| **Megasphaera A 38685** | 1.833 | 0.719 | 0.011 | 12% |
| **Faecalibacterium** | 2.448 | 1.198 | 0.041 | |
| **Sow treatment group:** | | | | |
| No treatment | | | | |
| Penicillin | -0.157 | 0.033 | <0.001 | |
| Tetracycline | -0.077 | 0.032 | 0.019 | |
| **Faecalibacterium** | 3.659 | 1.056 | 0.001 | 33% |

**Table S12.** Results from univariable general linear models evaluating associations between piglets' (n = 82) fecal microbiota relative abundace at the genus level at weaning and average daily weight gain (kg/day) from weaning to the end of the study period (ADG-nursing), both before and after adjusting for the sows' treatment group to assess potential confounding effects. All other potential confounders (sow parity, sow body contition score and pregnacy room) were included in the univariable models (results not shown). The genera selected for the univariable models did not show significant associations with weight gain in the multivariable model, but were significantly associated with the sows' treatment group.

| Variable | Coeff. | SE | p-value | Change of coeff. |
|---|---|---|---|---|
| **VUNA01** | 4.36 | 1.627 | 0.007 | |
| **Sow treatment group:** | | | | |
| No treatment | | | | |
| Penicillin | -0.112 | 0.037 | 0.003 | |
| Tetracycline | -0.061 | 0.036 | 0.09 | |
| **VUNA01** | 2.620 | 1.648 | 0.112 | 40% |
| **Alloprevotella (log2)** | 0.013 | 0.008 | 0.100 | |
| **Sow treatment group:** | | | | |
| No treatment | | | | |
| Penicillin | -0.123 | 0.036 | 0.001 | |
| Tetracycline | -0.063 | 0.037 | 0.086 | |
| **Alloprevotella (log2)** | 0.009 | 0.008 | 0.238 | 30% |

**Table S13.** Results from a multivariable general linear model evaluating associations between piglets' (n = 82) fecal microbiota Shannon index at weaning and average daily weight gain (kg/day) from weaning to the end of the study period (ADG-nursing), both before and after including the sow treatment group to assess potential confounding effect.

| Variable | Coeff. | SE | p-value | Wald test p-value |
|---|---|---|---|---|
| **Sow parity group:**[1] 2-8 | | | | <0.001 |
| **Sow body condition score group:**[1] 2.5, 3, 3.5 and 4 | | | | <0.001 |
| **Sow pregnancy room:** | | | | |
| 1 | 0 | | | |
| 2 | 0.003 | 0.036 | 0.936* | |
| **Sex:** | | | | |
| Female | 0 | | | |
| Male | -0.029 | 0.017 | 0.096* | |
| Shannon | 0.037 | 0.017 | 0.038 | |
| **Intercept** | 0.497 | 0.096 | <0.001 | |
| **ADG-nursing and shannon index association after icluding sow treatement group** | | | | **Change of coeff.** |
| **Sow treatment group:** | | | | |
| No treatment | 0 | | | |
| Penicillin | -0.126 | 0.034 | <0.001 | |
| Tetracycline | -0.059 | 0.035 | 0.086 | |
| Shannon | 0.036 | 0.016 | 0.029 | 2% |
| **Intercept** | 0.391 | 0.093 | <0.001 | |

[1] Group differeces are omitted from the table
* Variable is included to control the confounding effect

**Table S14.** Results from a multivariable general linear model evaluating association between piglets' (n = 76) fecal microbiota Shannon index at the end of the study period and average daily weight gain (kg/day) from weaning to the end of the study period (ADG-nursing), both before and after including the sow treatment group to assess potential confounding effect.

| **Variable** | **Coeff.** | **SE** | **p-value** | **Wald test p-value** |
|---|---|---|---|---|
| **Sow parity group:**[1] 2-8 | | | | <0.001 |
| **Sow body condition score group:**[1] 2.5, 3, 3.5 and 4 | | | | <0.001 |
| **Sow pregnancy room:** | | | | |
| 1 | 0 | | | |
| 2 | 0.037 | 0.041 | 0.351* | |
| Shannon | -0.064 | 0.029 | 0.029 | |
| **Intercept** | 0.898 | 0.128 | <0.001 | |
| **ADG-nursing and shannon index association after icluding sow treatement group** | | | | **Change of coeff.** |
| **Sow treatment group:** | | | | |
| No treatment | 0 | | | |
| Penicillin | -0.124 | 0.034 | <0.001 | |
| Tetracycline | -0.058 | 0.034 | 0.092 | 17% |
| Shannon | -0.053 | 0.026 | 0.049 | |
| **Intercept** | 0.747 | 0.127 | <0.001 | |

[1] Group differeces are omitted from the table
* Variable is included to control the confounding effect
