## Supplementary figures and images for "Impact of gestational antibiotics on maternal and offspring gut microbiota and growth in pigs"

### Supplementary Figure S1

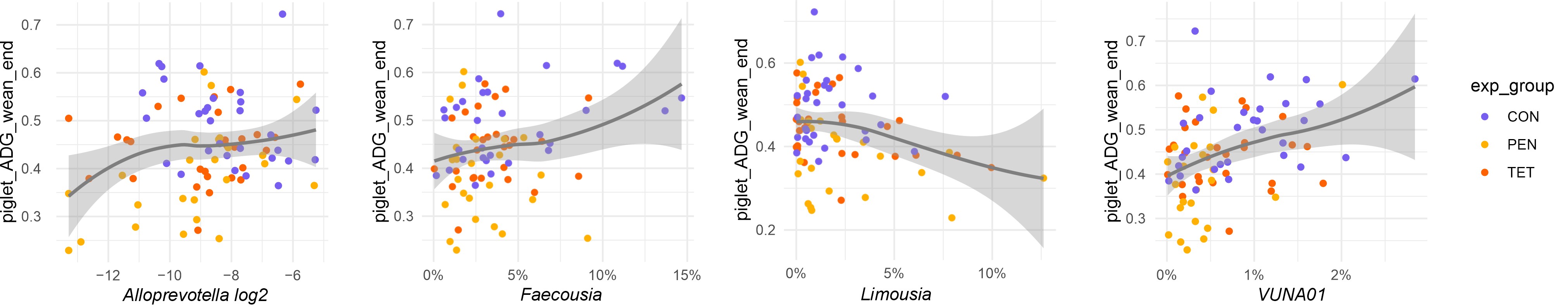
